# Reductive TCA cycle enzymes and reductive amino acid synthesis pathways contribute to electron balance in a *Rhodospirillum rubrum* Calvin cycle mutant

**DOI:** 10.1101/614065

**Authors:** Alexandra L. McCully, Maureen C. Onyeziri, Breah LaSarre, Jennifer R. Gliessman, James B. McKinlay

**Author notes:** Equal contributions (Alexandra L. McCully, Maureen C. Onyeziri).

## Abstract

Purple nonsulfur bacteria (PNSB) use light for energy and organic substrates for carbon and electrons when growing photoheterotrophically. This lifestyle generates more reduced electron carriers than are required for biosynthesis, even during consumption of some of the most oxidized organic substrates like malate and fumarate. Excess reduced electron carriers must be oxidized for photoheterotrophic growth to occur. Diverse PNSB commonly rely on the CO_2_-fixing Calvin cycle to oxidize excess reduced electron carriers. Some PNSB also produce H_2_ or reduce terminal electron acceptors as alternatives to the Calvin cycle. *Rhodospirillum rubrum* Calvin cycle mutants defy this trend by growing phototrophically on malate or fumarate without H_2_ production or access to terminal electron acceptors. We used ^13^C-tracer experiments to examine how a *Rs. rubrum* Calvin cycle mutant maintains electron balance under such conditions. We detected the reversal of some TCA cycle enzymes, which carried reductive flux from malate or fumarate to α-ketoglutarate. This pathway and the reductive synthesis of amino acids derived from α-ketoglutarate are likely important for electron balance, as supplementing the growth medium with α-ketoglutarate-derived amino acids prevented *Rs. rubrum* Calvin cycle mutant growth unless a terminal electron acceptor was provided. Flux estimates also suggested that the Calvin cycle mutant preferentially synthesized isoleucine using the reductive threonine-dependent pathway instead of the less-reductive citramalate-dependent pathway. Collectively, our results suggest that alternative biosynthetic pathways can contribute to electron balance within the constraints of a relatively constant biomass composition.

## Introduction

Purple nonsulfur bacteria (PNSB) are a metabolically versatile group that are often cultured under anaerobic conditions that support photoheterotrophic growth, wherein light is used for energy and organic compounds are used for carbon and electrons. This lifestyle presents a challenge in maintaining electron balance, as oxidative metabolic pathways generate an excess of reduced electron carriers (i.e., reductant) (1, 2). Oxidation of excess reductant is essential for growth and is often coupled to CO_2_ fixation in the Calvin cycle. This is true even for growth on substrates that are more oxidized than the average composition of cellular biomass, such as acetate, succinate, malate, and fumarate (Table 1) (3). Calvin cycle mutants are generally incapable of photoheterotrophic growth unless another means of electron disposal is possible, such as the reduction of terminal electron acceptors, like dimethylsulfoxide (DMSO) (4-7), or H_2_ production via nitrogenase (3, 8-11). Such trends have been observed for several PNSB including *Rhodobacter sphaeroides* (4, 5, 7, 11), *Rhodobacter capsulatus* (6, 12), and *Rhodopseudomonas palustris* (3, 8-10). An exception to this general observation is *Rhodospirillum rubrum*, for which Calvin cycle mutants grow photoheterotrophically on malate and fumarate without access to terminal electron acceptors and without producing H_2_ (9, 13). However, a terminal electron acceptor was still required by *Rs. rubrum* Calvin cycle mutants for phototrophic growth on succinate, a more reduced substrate (Table 1) (9). The mechanism by which *Rs. rubrum* Calvin cycle mutants maintain electron balance during phototrophic growth on malate or fumarate is unknown, but the fact that growth only occurs on these relatively oxidized substrates suggests that there is an inherent constraint on the amount of excess reducing equivalents that can be dealt with.

**Table 1.** Oxidation states of substrates mentioned in this study relative to biomass.

| Substrate | Formula | Oxidation state<sup>a</sup> |
| --- | --- | --- |
| carbon dioxide | CO<sub>2</sub> | +4 |
| formate | CH<sub>2</sub>O<sub>2</sub> | +2 |
| fumarate | C<sub>4</sub>H<sub>4</sub>O<sub>4</sub> | +1 |
| malate | C<sub>4</sub>H<sub>6</sub>O<sub>5</sub> | +1 |
| succinate | C<sub>4</sub>H<sub>6</sub>O<sub>4</sub> | +0.5 |
| acetate | C<sub>2</sub>H<sub>4</sub>O<sub>2</sub> | 0 |
| biomass | CH<sub>1.8</sub>N<sub>0.18</sub>O<sub>0.38</sub> | -0.5 |

<sup>a</sup>Values were determined for each carbon atom as described previously (10, 14) and then averaged by dividing the sum by the number of carbon atoms. Formulas for the acids of organic acids are shown.
<sup>b</sup>Biomass composition of *R. palustris* 42OL was used (15).

Alternative central metabolic pathways that could maintain electron balance during phototrophic growth in the absence of the Calvin cycle have been explored in silico (16), and some have been ruled out. For example, phototrophic growth by Calvin cycle mutants on malate could be possible if the cells produced formate (16). However, we previously found that *Rs. rubrum* Calvin cycle mutants did not excrete formate, nor did they accumulate polyhydroxybutyrate, an electron rich storage polymer that could potentially help maintain electron balance (9). There is also evidence that an alternative CO_2_-fixing pathway could be involved, as an *Rs. rubrum* Calvin cycle mutant was observed to grow when CO_2_ was the sole carbon source (17). In *Rb. sphaeroides* and *Rs. rubrum*, the CO_2_-fixing ethylmalonyl-CoA pathway can substitute for the Calvin cycle to achieve electron balance during phototrophic growth on acetate (18, 19). However, during phototrophic growth on malate, the ethylmalonyl-CoA pathway is not expected to be active because malate is also the product of the pathway and such a cycle would therefore be oxidative overall (16). Another CO_2_-fixing mechanism that has been proposed to participate in electron balance in *Rs. rubrum* Calvin cycle mutants is the partial reversal of the TCA cycle, specifically, the reductive activity of α-ketoglutarate (αKG) synthase that combines succinyl-CoA + CO_2_ + reductant to form αKG (16, 20, 21).

Here we used ^13^C-tracer experiments to gain insight into electron-balancing mechanisms in a *Rs. rubrum* Calvin cycle mutant. Labeling patterns suggested net reductive TCA flux to αKG from fumarate and malate. The reductive TCA cycle branch feeds into canonical reductive pathways for the synthesis of αKG-derived amino acids. These amino acid synthesis pathways likely also contribute to electron balance in a *Rs. rubrum* Calvin cycle mutant, as supplementing the medium with αKG-derived amino acids prevented phototrophic growth on malate unless DMSO was provided as a terminal electron acceptor. ^13^C-tracer experiments using H_2_-producing strains of *Rp. palustris*, which requires the Calvin cycle or H_2_ production to grow phototrophically on malate or fumarate, suggest that both fumarate reductase and αKG synthase are required for growth on malate or fumarate without the Calvin cycle. Flux estimates also revealed the preferential use of the reductive threonine-dependent isoleucine synthesis pathway over the less-reductive citramalate-dependent pathway in a *Rs. rubrum* Calvin cycle mutant but not in *Rp. palustris*. Thus, *Rs. rubrum* appears to have flexibility within its biosynthetic pathways that *Rp. palustris* does not, which can be used to satisfy electron balance during phototrophic growth on substrates as oxidized as malate and fumarate.

## Results and Discussion

### ^13^C-metabolic flux analysis of *Rs. rubrum* strains

To gain insight into how *Rs. rubrum* maintains electron balance without the Calvin cycle during phototrophic growth on relatively oxidized substrates, we performed ^13^C-metabolic flux analysis (^13^C-MFA) to estimate in vivo metabolic fluxes. In ^13^C-MFA, the activities of different pathways generate signature patterns of ^12^C and ^13^C that are imprinted on proteinaceous amino acids, which can be deciphered using gas chromatography-mass spectrometry (GC-MS) (22). Software is then used to identify a set of fluxes that can explain the observed mass isotopomer distributions (MIDs; i.e., labeling patterns). Directly measured fluxes, such as those for excreted products and those needed to generate a defined biomass composition, are also taken into account. We grew either wild-type (WT) *Rs. rubrum* or a Calvin cycle mutant lacking genes for phosphoribulokinase (PRK) and ribulose-1,5-bisphosphate carboxylase (RuBisCO), hereon referred to as the ΔPRKΔRubisco mutant (13), in a defined medium with [1,4-^13^C]fumarate. Cells harvested in exponential phase were used to determine MIDs in proteinaceous amino acids. No H_2_ was detected at the time of harvesting, indicating that H_2_ production was not involved in electron balance. High performance liquid chromatography (HPLC) analysis of culture supernatants revealed that malate was excreted by both strains, but more so by the ΔPRKΔRubisco mutant (46.2 ± 24.9 [± SD] mol malate/100 mol fumarate] than the WT strain (8.5 ± 3.9 mol malate/100 mol fumarate). Malate excretion during growth on fumarate has previously been observed in other bacteria, including *Rp. palustris* (3) and *Geobacter sulfurreducens* (23). Malate accumulation could result from an imbalance between the activities of fumarase and oxidative malate dehydrogenase when fumarate concentrations exceed those normally encountered in nature. Alternatively, malate accumulation might be necessary to make malate oxidation thermodynamically feasible (23). The higher level of malate excretion in the ΔPRKΔRubisco mutant could be due to either, or both, a slower rate of electron carrier oxidation that limits the NAD^+^ available for malate oxidation or an accumulation of NADH that increases the thermodynamic barrier for malate oxidation.

We then used the ^13^C-MFA software INCA (24) to estimate a set of fluxes that could explain the observed amino acid MIDs (Table S1), the malate excretion levels, and the *Rs. rubrum* biomass composition (Table S2 and S3) using a redox-constrained model of *Rs. rubrum* central metabolism and amino acid synthesis pathways (Table S4). For the biomass compositions of the *Rs. rubrum* strains, we determined the protein content (75 ± 4% of the dry cell weight [DCW]; statistically similar for both strains) and the amino acid composition (Table S3), and we assumed that the remaining 25% of the DCW had a composition similar to that of *Rp. Palustris* (10). Our models originally included the ethylmalonyl-CoA pathway. Optimal solutions indicated that a small reverse flux (<1% of the fumarate uptake rate) or exchange flux through the ethylmalonyl-CoA pathway was needed to explain the level of m+2 glutamate (Glu) that we observed. However, using INCAs simulation tool, we also found that exchange flux through only one ethylmalonyl-CoA pathway reaction, namely, malate synthase, was sufficient to explain the observed level of m+2 Glu (Fig. S1). We considered that exchange flux through malate synthase was more likely than net flux in either direction through the entire 12-step pathway, which is expected to be upregulated only during growth on C2 substrates like acetate (25), and as described in the introduction, would not contribute to electron balance during growth on compounds like malate or fumarate (16). The flux solutions we describe hereafter therefore include reversible malate synthase but exclude the ethylmalonyl-CoA pathway. Optimal flux solutions showed good agreement between the simulated and measured MIDs (Fig. S2). There was also good agreement between most measured fluxes and simulated fluxes except for summed biosynthetic redox reactions in the ΔPRKΔRubisco mutant and malate excretion for both strains (Fig. S2). These few disagreements suggest that our flux solutions contain some quantitative errors but, nevertheless, should point to qualitative trends that could contribute to electron balance, which we address below.

### *Rs. rubrum* flux solutions suggest reductive flux from fumarate to αKG

The flux maps for both WT *Rs. rubrum* and the ΔPRKΔRubisco mutant revealed a bifurcation of the TCA cycle; specifically, flux through an oxidative arm and flux through a reductive arm converged at αKG (Fig. 1). Our flux estimates are consistent with the notion that *Rs. rubrum* lacks a complete reverse TCA cycle but could still operate a reductive branch involving αKG synthase (20, 21). Flux through αKG synthase was previously proposed as a mechanism by which a Calvin cycle mutant could achieve electron balance (16, 21), based on αKG synthase activity detected in *Rs. rubrum* cell extracts (20). We questioned whether the reductive flux from fumarate to αKG, however small the estimated magnitude (0.1 and 0.6 mole % of the fumarate uptake rate in the WT and ΔPRKΔRubisco strains, respectively), could be important for electron balance.

**Fig. 1.**
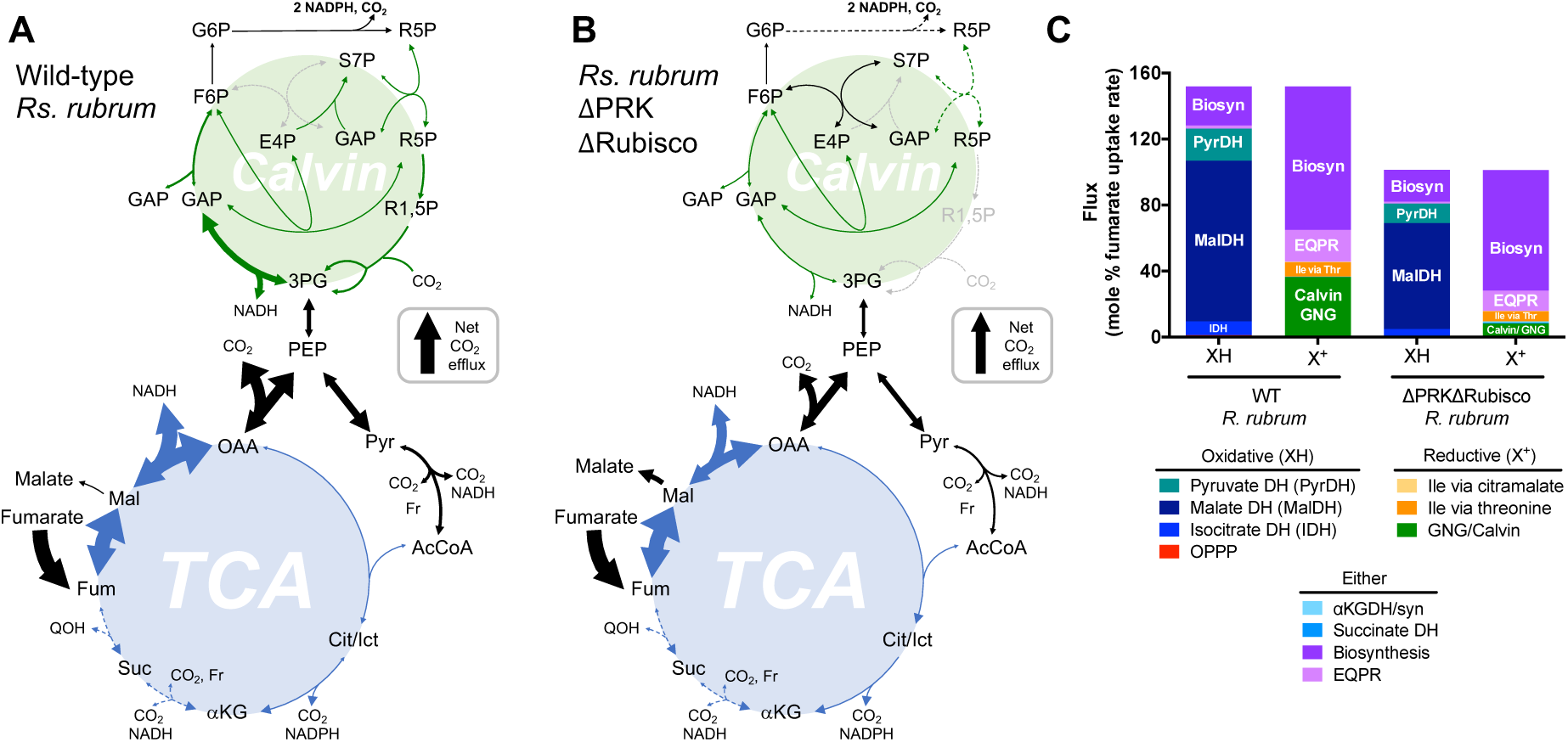
^13^C-tracer experiments suggest net reductive TCA flux to αKG in *Rs. rubrum* grown on fumarate. (A, B) Optimal flux solutions for WT *Rs. rubrum* (UR2) (A) and the ΔPRKΔRubisco mutant (UR2557) (B) grown phototrophically on [1,4-^13^C]fumarate. Flux magnitudes (mol % of the fumarate uptake rate) are indicated by solid arrow thickness. Dashed arrows indicate fluxes with magnitudes less than 1 mol % of the fumarate uptake rate. Net flux direction is indicated by an enlarged arrowhead. (C) Comparison of fluxes through redox reactions. XH columns refer to oxidative pathways that reduce electron carriers, whereas X^+^ columns refer to reductive pathways that oxidize electron carriers. EQPR is the synthesis of Glu, Gln, Pro, and Arg estimated from the total flux to αKG multiplied by the proportion of αKG used for each amino acid (Table S3) and then by the number of oxidation or reduction steps in each synthesis pathway. DH, dehydrogenase; OPPP, oxidative pentose phosphate pathway; aKGDH/syn, αKG DH/synthase; GNG/Calvin, glyceraldehyde-3-phosphate DH in either gluconeogenesis (GNG) or the Calvin cycle (Calvin). (A-C) Absolute flux values were 1.73 ± 0.09 and 0.66 ± 0.16 mmol fumarate/g DCW/h ± SD for the WT strain and the ΔPRKΔRubisco mutant, respectively. All flux values are in Table S4.

We first wanted to understand the labeling patterns that would be generated by reductive flux from fumarate to αKG. Higher proportions of m+2 and m+3 Glu were observed in the ΔPRKΔRubisco mutant compared to WT *Rs. rubrum* (Fig. 2). As noted above, malate synthase exchange flux combined with either reductive or oxidative flux to αKG could generate m+2 Glu (Fig. S1). From similar labeling experiments with *Rp. palustris*, we knew that m+3 metabolites, including m+3 Glu, can be a signature of Calvin cycle activity during growth on substrates with ^13^C-labeled carboxyl groups (3). Without the Calvin cycle, we reasoned that m+3 Glu could be generated from reductive flux from [1,4-^13^C]fumarate to αKG when ^13^CO_2_ is combined with [1,4-^13^C]succinyl-CoA to form [1,2,5-^13^C]αKG, from which Glu is derived (Fig. 2A). The ^13^CO_2_ needed for such a mechanism would come from the decarboxylation reactions stemming from substrates with ^13^C-labeled carboxyl groups (Fig. 2A) (3). Unlike m+3 Glu, other m+3 amino acids were less abundant in the ΔPRKΔRubisco mutant than in the WT strain, which can also generate m+3 metabolites via the Calvin cycle (Fig. 2A, Table S1). Thus, the mechanism by which m+3 Glu was generated in the ΔPRKΔRubisco mutant must be within a few reactions from Glu biosynthesis; reductive TCA flux from fumarate to αKG is consistent with such a mechanism. Adjusting malate synthase activity using INCAs simulation tool indicated that malate synthase cannot contribute to generating m+3 Glu, suggesting that reductive αKG synthase is solely responsible for generating m+3 Glu in the ΔPRKΔRubisco mutant.

**Fig. 2.**
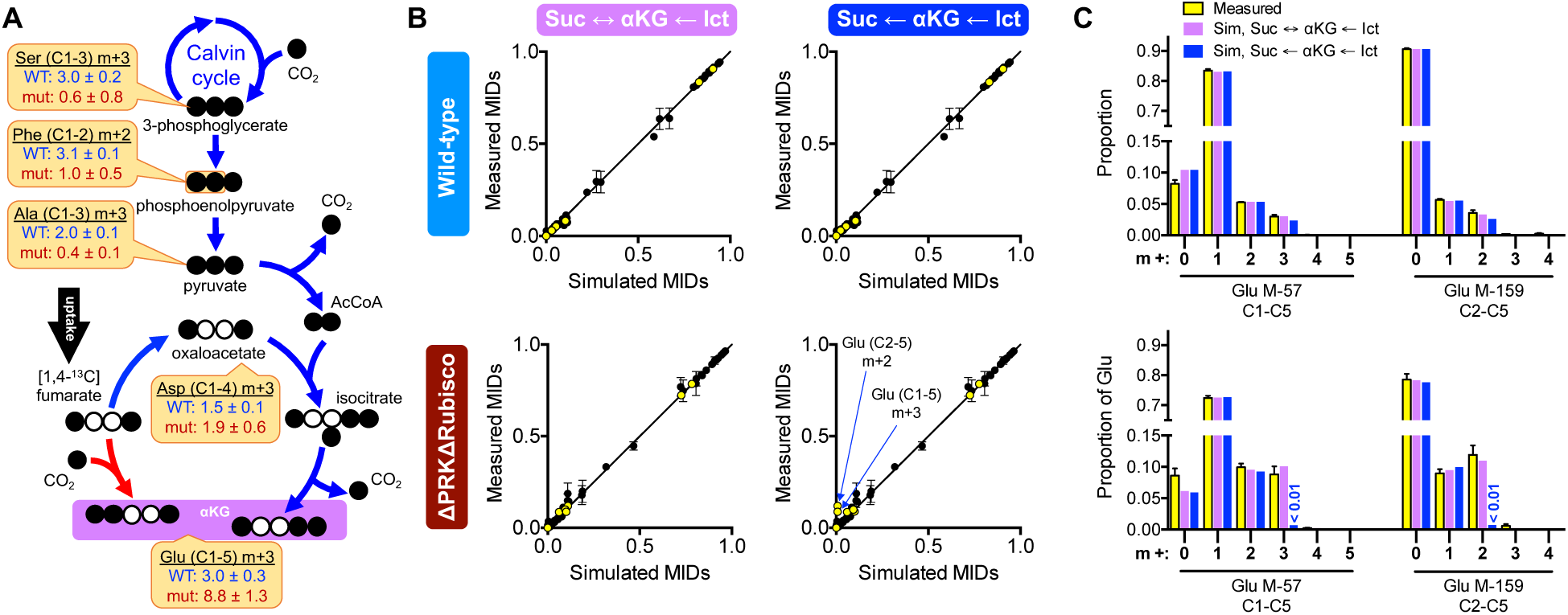
m+3 Glu is a signature of reductive flux from [1,4-^13^C]fumarate to αKG in the *Rs. rubrum* ΔPRKΔRubisco mutant. (A) m+3 Glu can be generated by both the Calvin cycle and reductive flux to αKG (red). However, the proportion of m+3 Asp is insufficient to explain the proportion of m+3 Glu by oxidative flux to αKG alone (blue). Yellow boxes show proportions for the indicated mass isotopomer ± SD. Decarboxylation reactions are the source of ^13^CO_2_ for carboxylation reactions. White, ^12^C; black, ^13^C. (B, C) Top, WT *Rs. rubrum* (UR2); bottom, *Rs. rubrum* ΔPRKΔRubisco (UR2557). (B) Linear regression of measured vs simulated MIDs from the optimal flux solutions for strains grown with [1,4-^13^C]fumarate. Simulations used a flux model that either allowed (left) or did not allow (right) for reversible flux between succinate (Suc) and αKG (αKG synthase). Irreversible flux from isocitrate (Ict) to αKG was assumed for all flux models. Yellow, Glu MIDs; black, MIDs for all other examined amino acids. (C) Comparison of measured and simulated (Sim) Glu MIDs from both models used in panel B. Error bars, SD; n = 4.

INCA’s simulation tool does not produce a new optimal solution every time an individual flux is adjusted. Thus, to confirm that reductive flux to αKG was the only mechanism that could generate m+3 Glu in the ΔPRKΔRubisco mutant, we ran fitting algorithms using a model in which αKG synthase flux was constrained to be strictly forward (i.e., oxidative). Whereas optimal solutions for WT *Rs. rubrum* models with strictly oxidative αKG synthase activity could fully explain the Glu labeling patterns, optimal solutions for the ΔPRKΔRubisco mutant resulted in a disagreement between the simulated and observed levels of m+3 Glu (Fig. 2B, C). In fact, without the Calvin cycle or reductive αKG synthase flux, the optimal solutions could not generate m+3 Glu (Fig. 2B, C). Thus, reductive flux to αKG is the only mechanism that can generate m+3 Glu in the ΔPRKΔRubisco mutant.

MIDs from a separate labeling experiment, wherein *Rs. rubrum* strains were cultured with unlabeled malate and NaH^13^CO_3_ as a source of ^13^CO_2_, also suggested net reductive flux to αKG for both WT and ΔPRKΔRubisco mutant strains (0.7 and 4.6 mole % of the fumarate uptake rate, respectively), in this case from malate (Fig. 3, yellow inset; Table S4). We expected that providing NaH^13^CO_3_ would result in the labeling of Glu by the reductive carboxylation of succinyl-CoA to αKG. However, both Glu and all other amino acids analyzed showed enrichment of m+1 isotopomers to varying degrees (Fig. 3; Table S1), which likely resulted from carboxylation via the Calvin cycle (WT strain only) and exchange flux through carboxylation reactions (both strains), such as those between PEP and oxaloacetate, between pyruvate and acetyl-CoA, and those in the citramalate-dependent isoleucine biosynthesis pathway; these possibilities were verified using INCAs simulation tool. Importantly, these reactions alone could not fully explain the measured Glu MIDs. In both strains, although only approximately 20% of aspartate, derived from oxaloacetate, had a mass of m+1, more than 30% of Glu, derived from αKG, had a mass of m+1 Glu. (Fig. 3). Thus, some of the m+1 Glu must have been derived from reductive flux from unlabeled malate to αKG. The ΔPRKΔRubisco mutant also exhibited a higher proportion of m+2 and m+3 Glu isotopomers compared to the WT strain (Fig. 3). These higher molecular weight Glu isotopomers could only be explained by the reductive carboxylation of single- and double-labeled succinyl-CoA, which itself must have been derived from exchange flux from single- and double-labeled oxaloacetate (Fig. 3). Both INCA simulations and sensitivity matrixes indicated that Glu labeling was relatively insensitive to malate synthase exchange flux under these experimental conditions, thus malate synthase was not a source of these labeling patterns. Thus, the results of two separate labeling experiments indicated reductive flux to αKG through multiple reactions typically associated with the reductive TCA cycle as a compensatory response to the absence of the Calvin cycle in the *Rs. rubrum* ΔPRKΔRubisco mutant.

**Fig. 3.**
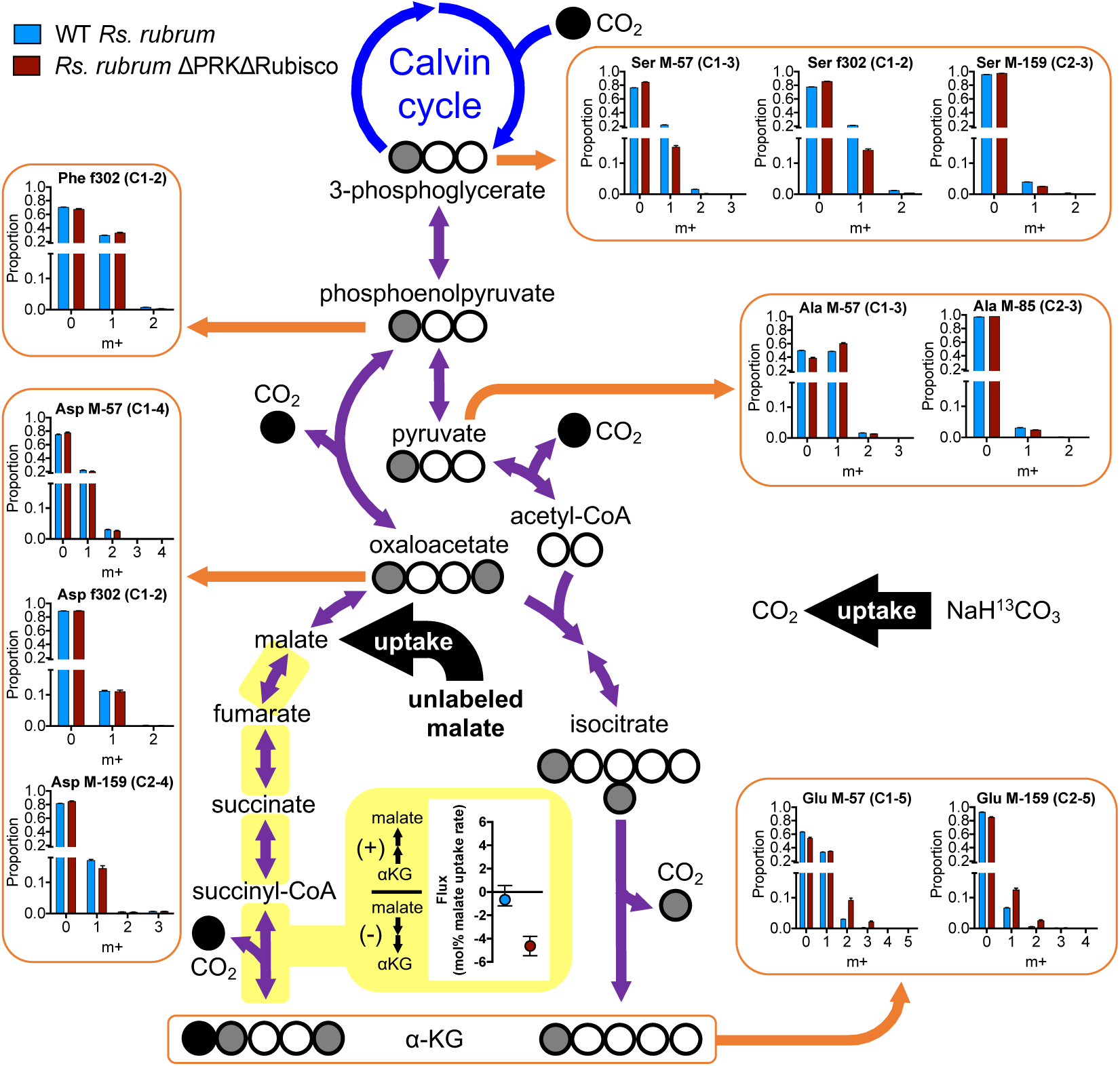
Flux estimates from experiments with unlabeled malate and NaH^13^CO_3_ suggest reductive net flux from malate to αKG in WT *Rs. rubrum* (UR2) and the ΔPRKΔRubisco mutant (UR2557). Only reductive flux to αKG can explain the high proportions of m+1, m+2, and m+3 Glu, all derived from αKG, relative to the corresponding MIDs of aspartate, derived from oxaloacetate. White, ^12^C; black, ^13^C; gray, partial labeling. Yellow inset: net flux value estimates between succinate and αKG (αKG synthase) with non-symmetrical upper and lower bounds determined by parameter continuation. Similar net flux values were obtained from malate to αKG (Table S4).

### αKG-derived amino acids inhibit growth of the *Rs. rubrum* ΔPRKΔRubisco mutant

Similar to our findings with photoheterotrophic *Rp. palustris* (3, 10), the relatively small flux to αKG in our *Rs. rubrum* flux maps was primarily dictated by the need for αKG as a biosynthetic precursor; although reductive to flux to αKG might play a role in electron balance, the flux did not exceed that needed to synthesize the amino acids Glu, Gln, Pro, and Arg. These four amino acids are all produced via reductive pathways and thus could also contribute to electron balance (Fig. 1C and 4A). In other bacteria, such as *E. coli*, the presence of each of these amino acids inhibits its own synthesis (26-30). We predicted that if these amino acid synthesis pathways are similarly regulated in *Rs. rubrum*, then their synthesis could be repressed by including them in the growth medium; in this manner, supplementation with these amino acids might prevent reductive flux to αKG. Addition of the four amino acids (EQPR) had no effect on the phototrophic growth trends of WT *Rs. rubrum* grown on malate but prevented ΔPRKΔRubisco mutant growth (Fig. 4B). These results could be interpreted to mean that adding Glu, Gln, Pro, and Arg inhibited the respective amino acid synthesis pathways and thereby prevented electron balance. An alternative explanation could be that, even at 0.2 mM each, the catabolism of these amino acids contributed to an excess of reductant. However, adding four amino acids derived from oxaloacetate (DTMK) did not affect growth of either WT or the ΔPRKΔRubisco mutant on malate (Fig. 4C). The addition of either 0.7 mM isoleucine (Fig. 5) or 0.7 mM leucine (data not shown), both of which should also generate reductant through their catabolism, also did not affect the growth of the ΔPRKΔRubisco mutant on malate. Thus, the inhibitory effect of Glu, Gln, Pro, and Arg on the ΔPRKΔRubisco mutant is specific to this amino acid mixture. To verify that the growth inhibition was linked to electron balance rather than to an unrelated form of toxicity, we repeated the growth experiment in medium supplemented with the terminal electron acceptor DMSO. Adding DMSO rescued ΔPRKΔRubisco mutant growth on malate in the presence of Glu, Gln, Pro, and Arg (Fig. 4D). Thus, the addition of Glu, Gln, Pro, and Arg created a growth-inhibiting electron imbalance in the ΔPRKΔRubisco mutant.

**Fig. 4.**
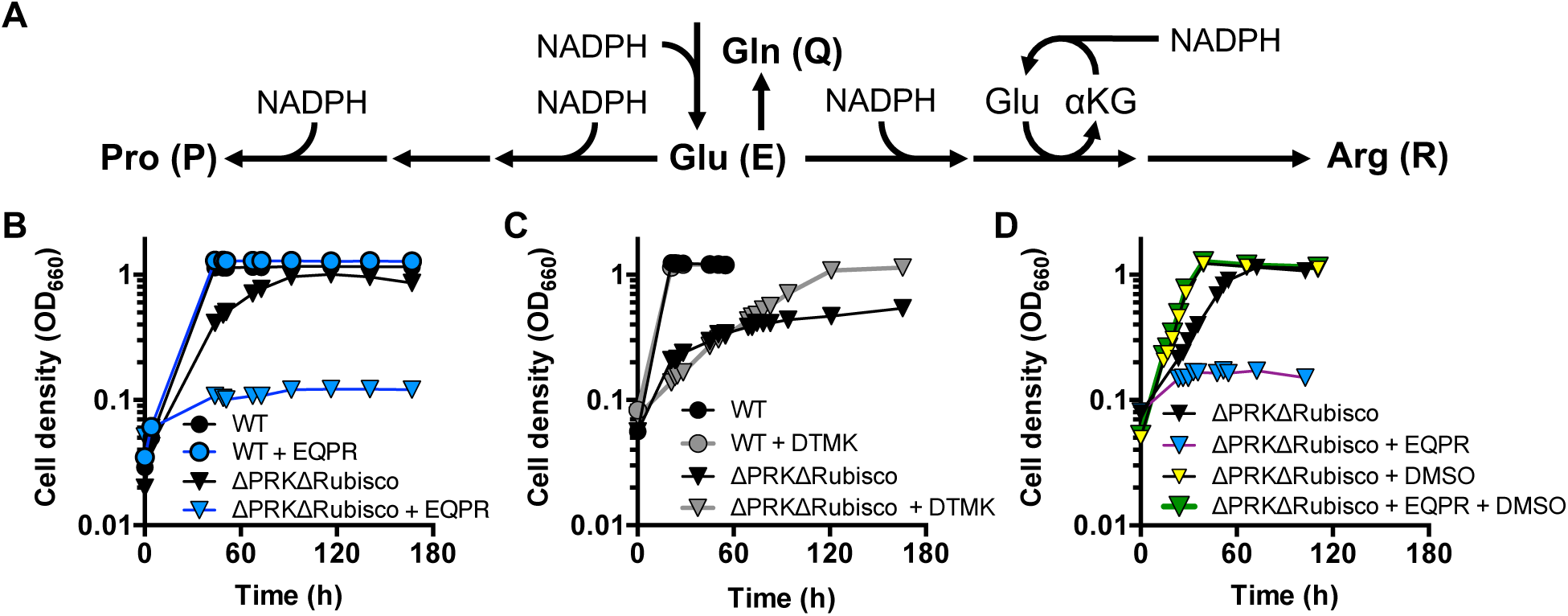
αKG-derived amino acids repress phototrophic growth of the *Rs. rubrum* ΔPRKΔRubisco mutant on malate. (A) Simplified pathway illustrating the redox reactions involved in the synthesis of the four amino acids synthesized from αKG. (B-D) Growth curves for WT *Rs. rubrum* and the ΔPRKΔRubisco mutant grown with a mixture of either glutamate, glutamine, proline, and arginine (EQPR) (B), with or without DMSO (D), or aspartate, threonine, methione, and lysine (DTMK) (C).

**Fig. 5.**
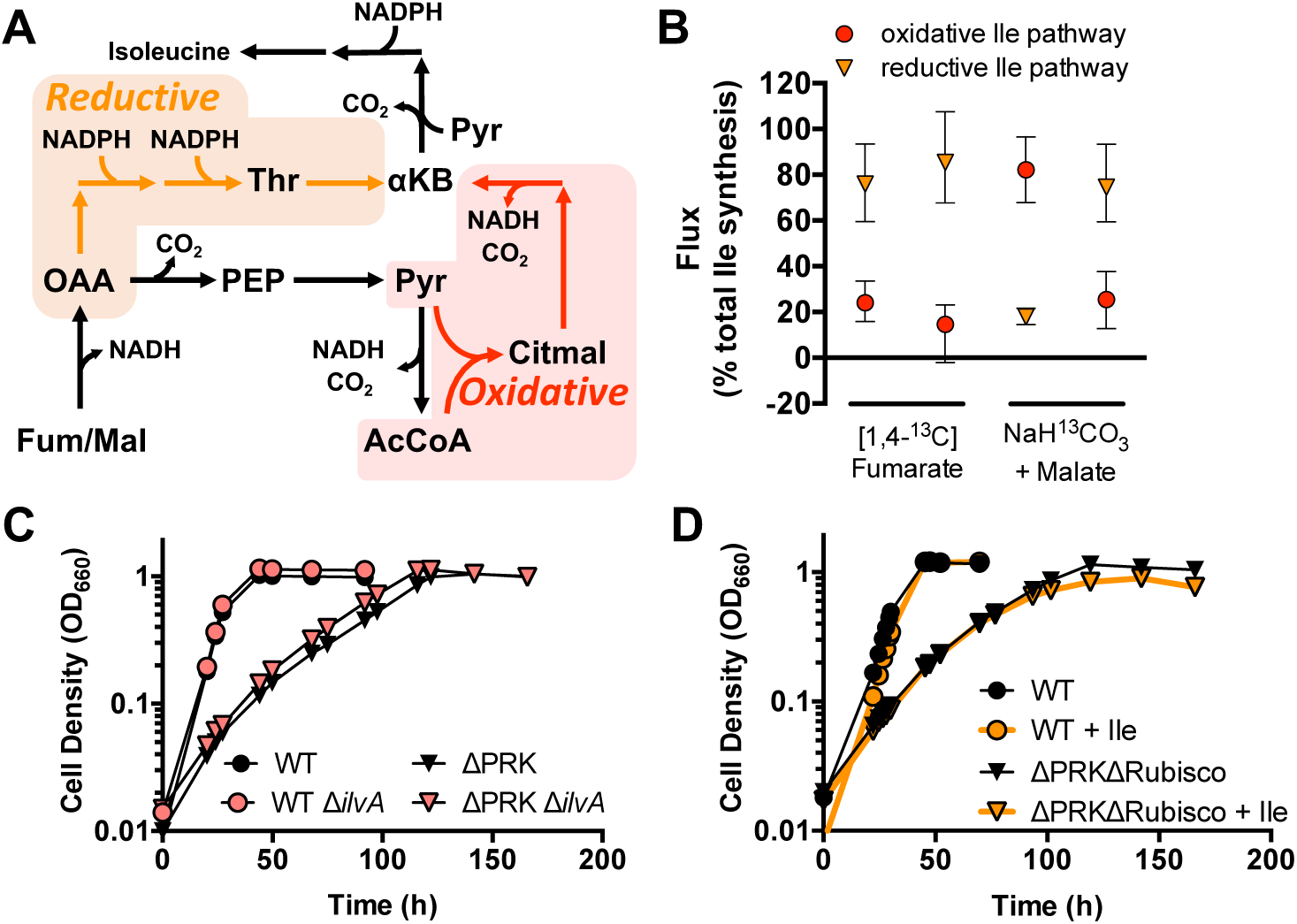
A non-essential reductive isoleucine synthesis pathway is favored in *Rs. rubrum* in the absence of the Calvin cycle. (A) *Rs. rubrum* has two pathways for isoleucine synthesis. The canonical threonine-dependent pathway (orange) is reductive, whereas the alternative citramalate-dependent pathway (red) is oxidative during growth on malate or fumarate. αKB, α-ketobutyrate. (B) Estimated fluxes as a percent of total isoleucine synthesis for the oxidative citramalate-dependent pathway and the reductive threonine-dependent pathway for the WT (UR2) and ΔPRKΔRubisco (UR2557) strains during growth on either [1,4-^13^C]fumarate or malate with NaH^13^CO_3_. All flux values are in Table S4. (C) Growth trends of WT and ΔPRK (UR2565) strains with or without deletion of *ilvA*, which encodes threonine deaminase in the reductive threonine-dependent pathway. (D) Growth trends of WT and ΔPRKΔRubisco strains with or without added isoleucine in the medium.

Our attempts to genetically determine whether reductive flux to αKG was essential were inconclusive. *Rs. rubrum* is annotated to have both irreversible αKG dehydrogenase (Rru_A1213–A1215) and reversible αKG synthase (Rru_A2720–A2722). We deleted Rru_A2721, annotated to encode the αKG synthase α-subunit, in a ΔPRK mutant (UR2565) (13). The ΔPRK strain cannot fix CO_2_ via the Calvin cycle, has similar growth trends to the ΔPRKΔRubisco mutant (9), and has the genetic background necessary for genetic engineering (unlike the ΔPRKΔRubisco mutant, the ΔPRK strain is not resistant to gentamycin, which is needed to select for integration of the suicide vector). Unfortunately, the ΔPRKΔRru_A2721 mutant exhibited a high incidence of H_2_ production (10 of 12 biological replicates across four independent experiments grew accompanied by H_2_ production), likely due to a spontaneous mutation that activates nitrogenase; such suppressor mutations are known to improve the growth of *Rs. rubrum* Calvin cycle mutants (31). The production of H_2_ could suggest that H_2_ production was necessary for the mutant to grow. However, the high frequency of H_2_ production prevented us from making a definitive conclusion about the essentiality of αKG synthase in the absence of the Calvin cycle.

### Isoleucine is produced primarily by a reductive pathway in the *Rs. rubrum* ΔPRKΔRubisco mutant

*Rs. rubrum* has two pathways for isoleucine synthesis. The canonical threonine-dependent pathway uses 1 oxaloacetate and 1 pyruvate, whereas the alternative citramalate-dependent pathway uses 1 AcCoA and 2 pyruvate (Fig. 5A) (32). Although both pathways are technically reductive, we herein consider the citramalate-dependent pathway to be oxidative because the path from fumarate or malate to 2 pyruvate + 1 AcCoA generates more reducing equivalents than can be oxidized in the subsequent synthesis of isoleucine (Fig. 5A). The optimal flux solutions for *Rs. rubrum* strains grown on [1,4-^13^C]fumarate estimated that the WT strain used the threonine-dependent pathway to make 76% of the isoleucine, whereas the ΔPRKΔRubisco mutant used this pathway to make 85% of the isoleucine (Fig. 5B). Flux solutions from conditions using unlabeled malate and Na^13^CO_3_ were more skewed, indicating that isoleucine was made primarily by the oxidative citramalate-dependent pathway in the WT strain, whereas the reductive threonine-dependent pathway was dominant in the ΔPRKΔRubisco mutant (Fig. 5B). Overall, the ^13^C-MFA solutions suggest that the threonine-dependent isoleucine synthesis pathway is used as a compensatory mechanism for maintaining electron balance in the absence of the Calvin cycle. The flexibility that the two isoleucine synthesis pathways impart in the management of electrons could explain why both seemingly redundant pathways are maintained in some bacteria (32-34).

To assess whether the reductive isoleucine pathway was essential for electron balance in the absence of the Calvin cycle, we deleted *ilvA* (Rru_A2877), encoding threonine dehydratase, in both WT *Rs. rubrum* and the ΔPRK mutant to prevent isoleucine synthesis via the threonine-dependent pathway. Separately, we also grew WT and ΔPRKΔRubisco *Rs. rubrum* strains in the presence of isoleucine, which has the potential to inhibit both isoleucine synthesis pathways if the pathways are regulated in *Rs. rubrum* as they are in *E. coli* (35) and *Leptospira interrogans* (36). Neither deleting *ilvA* (Fig. 5C) nor adding isoleucine to the growth medium (Fig. 5D) affected the growth trends of either the WT or the Calvin cycle mutants. No H_2_ accumulated during the course of these experiments. Thus, although the ^13^C labeling patterns suggested that the flux through the two isoleucine synthesis pathways was influenced by the need to maintain electron balance, the reductive threonine-dependent pathway is not essential for electron balance in *Rs. rubrum* Calvin cycle mutants.

### Sulfide production does not participate in electron balance in *Rs. Rubrum*

Reduction of sulfate to sulfide by a spontaneous mutant of *Rb. sphaeroides* was previously implicated as an alternative electron-balancing mechanism during photoheterotrophic growth (37). Thus, we also considered whether sulfate reduction could contribute to electron balance in *Rs. rubrum*. However, using lead acetate strips, we did not detect any sulfide in either WT or ΔPRKΔRubisco mutant cultures (Fig. S3).

### Why is the Calvin cycle required by *Rp. palustris* for photoheterotrophic growth?

*Rs. rubrum* Calvin cycle mutants can grow on fumarate and malate, whereas *Rp. palustris* Calvin cycle mutants cannot unless they can dispose of excess electrons as H_2_ (9). The *R. palustris* CGA009 genome is annotated to have an α-oxoacid ferredoxin oxidoreductase (RPA1225– 1228), suggesting that it could be capable of reductive αKG synthase activity. We considered that the reason *Rp. palustris* Calvin cycle mutants cannot grow on malate or fumarate might be due to an inability to reduce fumarate, which would deny αKG synthase from participating in electron balance during growth on fumarate and malate. All *Rp. palustris* strains tested by other groups have not exhibited fumarate reductase activity, unlike *Rs. rubrum* (38, 39). In *Rs. rubrum*, fumarate reductase activity is dependent on rhodoquinone (38), a compound that is notably absent from all *Rp. palustris* strains that have been analyzed (38, 40-42). Our BLAST searches against the *R. palustris* CGA009 genome (43) using the only known *Rs. rubrum* protein required for rhodoquinone synthesis, namely, the methyltransferase RquA, (Rru_A3227) (44), also suggested that *Rp. palustris* cannot synthesize rhodoquinone; the top hit was RPA0138, annotated as the chemotaxis methyl transferase CheR1, with only 31% coverage and 28% amino acid identity (E value: 0.002).

To assess whether *R. palustris* is capable of fumarate reductase and reductive αKG synthase activities in vivo, we performed four separate labeling experiments with an H_2_-producing *Rp. palustris* NifA* strain, CGA676, and its ΔPRKΔRubisco mutant counterpart, CGA4011 (9), and either [1,4-^13^C]succinate, [1,4-^13^C]fumarate, unlabeled succinate with NaH^13^CO_3_, or unlabeled malate with NaH^13^CO_3_. We used two models for each strain: one that allowed for reversible αKG synthase activity, and another that allowed for only oxidative αKG synthase activity. If *Rp. palustris* carried out reversible αKG synthase activity under these growth conditions then we expected that reductive αKG synthase activity would be required to explain Glu labeling patterns in the NifA* ΔPRKΔRubisco mutant during growth on succinate. If *Rp. palustris* is incapable of fumarate reductase activity then we expected that reductive αKG synthase activity would not be required to explain Glu labeling patterns on fumarate or malate, as there would be no way to reduce either of these compounds to the succinyl-CoA needed for reductive αKG synthase activity.

In models that allowed for reversible of αKG synthase activity, the optimal solutions showed net reductive flux to αKG when the NifA*ΔPRKΔRubisco mutant was grown with either [1,4-^13^C]succinate or succinate with NaH^13^CO_3_ (Fig. 6A). Net oxidative αKG synthase flux was estimated in all *Rp. palustris* NifA* conditions and in NifA*ΔPRKΔRubisco mutant conditions supplied with either [1,4-^13^C]fumarate or malate with NaH^13^CO_3_ (Fig. 6A). These solutions support the notion that *Rp. palustris* is capable of reductive αKG synthase activity but not fumarate reductase activity. The estimated reductive αKG synthase flux in the NifA*ΔPRKΔRubisco strain only could suggest that reductive αKG synthase activity provided some compensatory electron-balancing activity during growth on succinate in the absence of the Calvin cycle, despite being unessential given the requirement for H_2_ production (3, 10). However, it is noteworthy that markedly less m+3 Glu was generated by *Rp. palustris* NifA* ΔPRKΔRubisco compared to *Rp. palustris* NifA* during growth on [1,4-^13^C]succinate (Fig. 6B). Similarly, the abundance of m+1, m+2, and m+3 Glu was less in *Rp. palustris* NifA* ΔPRKΔRubisco compared to *Rp. palustris* NifA* when grown with succinate with NaH^13^CO_3_. These observations are opposite of the trends observed with *Rs. rubrum* and thus strongly suggest that reductive flux to αKG is not a general mechanism used by *Rp. palustris* in response to the loss of the Calvin cycle, at least during phototrophic growth on succinate when H_2_ is produced.

**Fig. 6.**
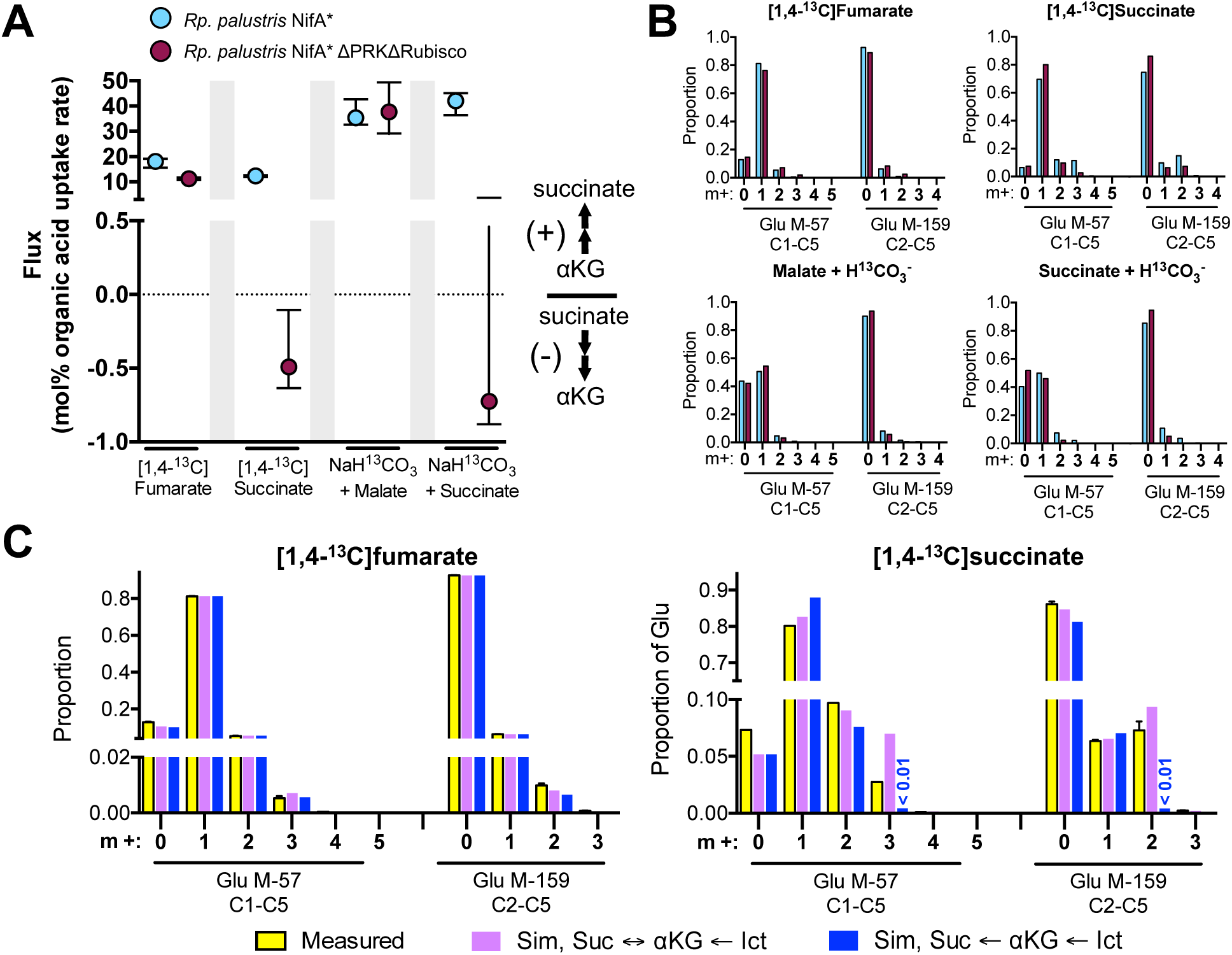
^13^C-labeling experiments suggest that *Rp. palustris* is capable of reductive αKG synthase activity but not fumarate reductase activity. (A, B) Net αKG synthase flux values (A) and Glu MIDs (B) from optimal flux solutions for *Rp. palustris* NifA* (CGA676) and *Rp. palustris* NifA* ΔPRKΔRubisco (CGA4011) grown phototrophically on various labeled substrates. (A) Flux values are mole % of either the fumarate, malate, or succinate uptake rate. Gray bars serve to separate experimental results. All flux values are in Table S4. (B) NifA* Glu MIDs were reported previously (3, 10). (C) Comparison of measured and simulated (Sim) Glu MIDs from models for *Rp. palustris* NifA* ΔPRKΔRubisco. Simulations either allowed or did not allow for reversible flux between succinate and αKG (αKG synthase). Error bars, SD; n = 3.

When the models were constrained to allow for only oxidative αKG synthase activity, Glu MIDs could still be explained for both *Rp. palustris* strains under most conditions. Experiments with NaH^13^CO_3_ did not determine αKG synthase activity with confidence in the NifA* ΔPRKΔRubisco mutant (Fig. 6A). However, similar to our observations with *Rs. rubrum*, reductive αKG synthase activity was required to generate m+3 Glu in the model for the NifA* ΔPRKΔRubisco mutant grown on [1,4-^13^C]succinate (Fig. 6C). The fact that reductive αKG synthase activity was required in this case but was not required to explain Glu labeling patterns when the NifA* ΔPRKΔRubisco mutant was grown on [1,4-^13^C]fumarate (Fig. 6C; data for other conditions not shown) supports the notion that there is no fumarate reductase activity to supply double-labeled succinate from which m+3 Glu would ultimately be derived (Fig. 2A). Thus, overall, there is good evidence that the inability to reduce fumarate is the reason why *Rp. palustris* Calvin cycle mutants cannot grow phototrophically on malate or fumarate without producing H_2_.

Although not expected to be an essential electron-balancing mechanism based on our data with *Rs. rubrum* (Fig. 8), we also examined whether H_2_-producing *Rp. palustris* adjusted fluxes through the same two isoleucine synthesis pathways (3, 10) in response to the loss of the Calvin cycle. Unlike for *Rs. rubrum*, the oxidative citramalate-dependent pathway was estimated to dominate under all but one condition, and flux through the reductive threonine-dependent pathway did not increase in response to the absence of the Calvin cycle (Fig. S4).

### Conclusion

Our data support previous results from our own group (9) and others (13) that *Rs. rubrum* can maintain electron balance during photoheterotrophic growth on malate and fumarate without the Calvin cycle, H_2_ production, or access to terminal electron acceptors. Our results suggest that a *Rs. rubrum* Calvin cycle mutant can maintain electron balance during phototrophic growth on malate and fumarate by selectively utilizing various biosynthetic pathways. The relatively static nature of biomass composition for a given growth condition places constraints on the extent to which different biosynthetic pathways can be employed to achieve electron balance and thus likely explains why a *Rs. rubrum* Calvin cycle mutant cannot grow with succinate, which has only two more electrons than malate or fumarate (Table 1). The two biosynthetic pathways we identified to contribute to electron balance in *Rs. rubrum* were (i) reductive flux from fumarate to αKG linked to the reductive synthesis of amino acids from αKG and (ii) the preferential use of the reductive threonine-dependent pathway for isoleucine synthesis instead of the less-reductive citramalate-dependent pathway. Given the relatively minor influence the pathways are estimated to have on electron balance (Fig. 1C), it is possible that other biosynthetic pathways not be detected by our approach, such as altered fatty acid composition, could also offer limited flexibility to contribute to electron balance. Flexibility in electron management could explain why some bacteria maintain seemingly redundant biosynthetic pathways, such as those for isoleucine synthesis and those supplying αKG for amino acid synthesis. Our flux estimates suggest that αKG can be synthesized by via both reduction and oxidation of TCA cycle intermediates from malate and fumarate in *Rs. rubrum* and from succinate in *Rp. palustris*. Such activities would add these two PNSB to a short list of bacteria known to produce αKG-derived amino acids via two separate pathways (45, 46). The previous detection of fumarate reductase activity in diverse PNSB (38, 39) raises the possibility that such dual pathways might be found in other PNSB as well and perhaps play a conserved role in electron balance during photoheterotrophic growth in these bacteria.

## Methods

### Strains and growth conditions

*Rs. rubrum* strains ΔPRKΔRubisco (UR2557, Δ*cbbM::*Gm^R^ Δ*cbbP*::Km^R^ (13)) and ΔPRK (UR2565, Δ*cbbP*::Km^R^ (13)) were derived from type-strain UR2 (47). *Rp. palustris* strains CGA676 (NifA*, (10)) and CGA4011 (NifA*, Δ*cbbLS*, Δ*cbbM*, Δ*cbbP*::Km^R^ (9)) were derived from type-strain CGA009. *Rs. rubrum* or *Rp. palustris* strains were revived from frozen stocks containing 10% DMSO or 25% glycerol, respectively, by streaking for single colonies on defined photosynthetic medium (PM) agar (48) supplemented with 10 mM disodium succinate and 0.1% yeast extract without antibiotics. Colonies were then used to inoculate 3 ml aerobic starter cultures of PM (*Rp. palustris*) or defined MA medium (*Rs. rubrum*) (9), each with 10 mM succinate. Aerobic starter cultures were grown at 30°C with shaking at 150 rpm. A 0.1-ml inoculum from stationary phase aerobic starter cultures was then transferred into 10 ml of anaerobic PM (*Rp. palustris*) or MA (*Rs. rubrum*) supplemented with 10 mM of an organic substrate, as indicated, in 28-ml anaerobic test tubes sealed with rubber stoppers under an argon headspace, as described (9). Where indicated, media were supplemented with specific amino acids (20 mg/L each), DMSO (60mM), or NaHCO_3_ (20 mM), prior to inoculation. Antibiotics were not used in comparative growth experiments or in ^13^C-labeling experiments but were otherwise added to media where appropriate. Gentamycin and kanamycin were each used at 100 µg/ml for *Rp. palustris* and at 10 µg/ml for *Rs. rubrum. Bacillus subtilis* 3610 (49) and *Escherichia coli* strains NEB10β (New England Biolabs) and WM3064 were cultured aerobically on LB agar or in LB broth. WM3064 is a diaminopimelic acid (DAP) auxotroph and was thus supplemented with 0.6 mM DAP (W. Metcalf, unpublished data). Gentamycin was used at 15 µg/ml for *E. coli*.

### Analytical techniques

Cell density was assayed by optical density (OD_660_) using a Genesys 20 visible spectrophotometer (Thermo-Fisher, Pittsburgh, PA). Specific growth rates were determined using measurements with values between 0.1 – 0.8 OD_660_, where a linear relationship between cell density and OD_660_ was maintained. H_2_ was sampled from culture headspace using a gas-tight syringe and analyzed using a Shimadzu GC-2014 gas chromatograph as described (50). Organic acid levels in culture supernatants were determined by HPLC (Shimadzu) as described (51). Sulfide was detected using lead acetate strips suspended above the cultures. Strips were removed and photographed once cultures reached stationary phase (52).

### *Rs. rubrum* biomass composition

Measurements were performed for the WT strain (UR2) and the ΔPRKΔRubisco mutant (UR2557) grown in MA medium. Culture samples were taken between 0.08 and 0.85 OD_660_ as measured using plastic cuvettes with a 1-cm path length. DCWs were determined as described (53) by filtering 30-mL of culture through preweighed 0.22-µm HA filters (Millipore) and then drying the filters overnight at 80 °C. Protein was quantified using a bicinchoninic acid assay (Pierce) as described (10). Amino acid composition was determined using a Hitachi L-8800 amino acid analyzer at the University of California, Davis Molecular Structure Facility. The polyhydroxybutyrate content was previously found to be below the detection limit (9). Because protein accounted for 75% of the DCW, the remaining DCW not accounted for in protein was assumed to comprise other major macromolecules at the same proportions as found in *Rp. palustris* and with the same monomeric composition. A common biomass composition was used for both strains because the slopes for linear regression of DCW/L vs. OD_660_ (400 ± 40 mg/L/ OD_660_) and protein vs. DCW (75 ± 4% of DCW) were statistically similar (p > 0.1, determined using the linear regression analysis function in Graphpad Prism 6.0).

### ^13^C-labeling experiments and metabolic flux analysis

Three or four biological replicates of each strain were inoculated from unlabeled starter cultures into media containing the indicated ^13^C-labeled substrate, provided at 100%. Cultures were harvested in exponential phase between 0.2 – 0.7 OD_660_ as described previously (10). Amino acids were obtained from the cell pellets by hydrolysis in 6 N HCl, the amino acids were derivatized, and then MIDs were determined using GC-MS as described (53) at the Indiana University Mass Spectrometry Facility. MIDs were corrected for natural abundances of all atoms except for carbons in amino acid backbones using the IsoCor software (54). Corrected amino acid MIDs and flux measurements (i.e., excreted malate or fumarate and fluxes derived from the biomass composition [Table S2]) were used with a metabolic model and the software INCA (24) to map the carbon atom transitions through the central metabolism of *Rs. rubrum* or *Rp. palustris* (Table S4). Our *Rs. rubrum* model included the TCA cycle, gluconeogenesis/glycolysis, the Calvin cycle, the pentose phosphate pathway, and the ethylmalonyl-CoA pathway (Table S4). Models that required that the sum of fluxes through reduction reactions equal the sum of fluxes through oxidation reactions included terms representing oxidized electron carriers (X) and reduced electron carriers (XH) in each redox reaction. Equations were also included to reflect the reductant required for biosynthesis and the reductant generated by biosynthesis. For each condition or model variation tested, at least 100 different arrangements of starting free parameter values were used. An optimal solution was selected based on that which had the lowest sum of squared residuals between the simulated and measured data sets for MIDs and biosynthetic and extracellular fluxes. Upper and lower bounds for flux values were determined using INCAs parameter continuation tool.

### *Rs. rubrum* strain construction

To create an in-frame deletion of Rru_A2721 (αKG synthase α-subunit), the region upstream of Rru_A2721 was amplified using primers BL501 (gacttctagactataacgatctggccgattggc) and BL524 (gactggtacccgtcatcatccgtctttcagagaag), and the region downstream was amplified using primers BL525 (gactggtacccagggagagtgatcatggatacg) and BL504 (gacttctagaactgggcttcgatttcggtgaa). The resulting PCR products were digested using KpnI and ligated together, and then the ligation reaction was used as template for a second round of PCR using primers BL501 and BL504. The resulting PCR product was purified, digested with XbaI, and then ligated into XbaI-digested pJQ200SK (55). To create an in-frame deletion of Rru_A2877 (*ilvA*, threonine deaminase), the upstream region was amplified using primers BL646 (gctggagctccaccgtgactccgtctggccccaag) and BL635 (tccccggcggtcatcgtccgccctcc), and the downstream region was amplified using primers BL636 (cgatgaccgccggggattagggcaaatc) and BL647 (attcctgcagcccggcgtcgattacaccggcctc). The upstream and downstream PCR products were mixed with pJQ200SK, which was PCR-linearized using primers BL644 (cggtggagctccagcttttg) and BL645 (ccgggctgcaggaattcg), and then assembled by Gibson Assembly (NEB). Ligation and Gibson reactions were transformed into *E. coli* NEB10β (New England Biolabs), and transformants were verified by PCR and sequencing. Vectors were first introduced into electrocompetent *E. coli* WM3064 and were then transferred to *Rs. rubrum* strains by conjugation on PM agar with yeast extract, succinate, and DAP. Mating patches were streaked for single recombinants on PM agar with yeast extract, succinate, and Gm (no DAP). Colonies were then grown in liquid medium without Gm to allow for a second recombination event. Cultures were plated on PM agar with yeast extract, succinate, and 10% sucrose, resultant colonies were screened for Gm sensitivity, and then Gm-sensitive colonies were screened by PCR for the desired deletions.

## Author statements

JBM conceived the study. All authors performed experiments. ALM, MCO, JRG, and JBM analyzed the data. JBM wrote the initial manuscript draft. ALM, BL, and JBM edited subsequent drafts. All authors reviewed and approved the final draft.

The authors declare no conflict of interest.

This work was funded by the Indiana University College of Arts and Sciences.

We thank GC Gordon for contributions to preliminary studies, JA Karty for GC-MS training and advice, and anonymous reviewers for insightful questions that led to the improvement of this work.

## Supplementary figures and legends

**Fig. S1.**
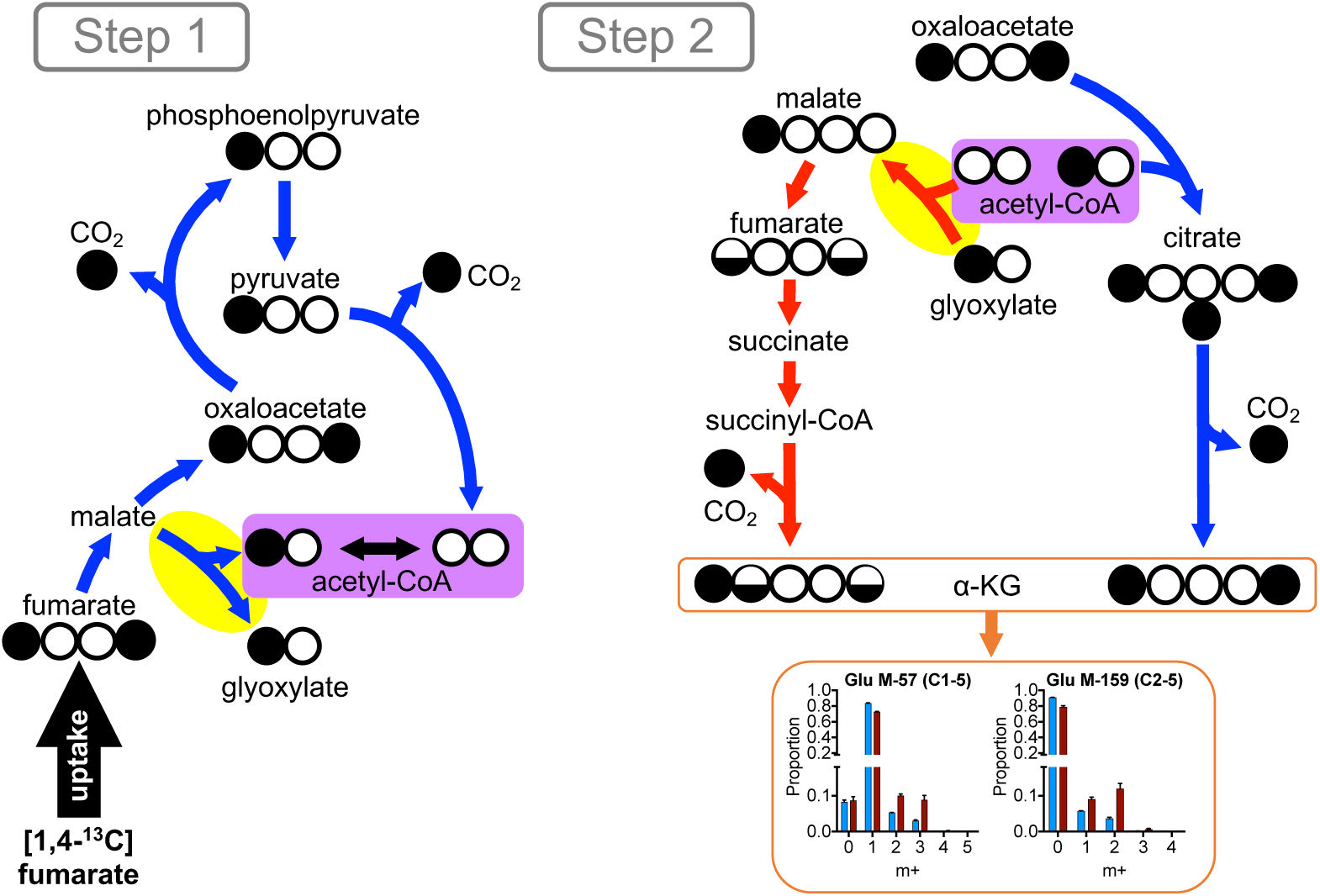
Exchange flux through malate synthase (highlighted yellow) can contribute to the generation of m+2 Glu in *Rs. rubrum* during growth on [1,4-^13^C]fumarate. Step 1. Reverse malate synthase activity generates single-labeled acetyl-CoA while pyruvate dehydrogenase generates unlabeled acetyl-CoA. Step 2. Forward malate synthase activity combines unlabeled acetyl-CoA with single-labeled glyoxylate to form single-labeled malate. Single-labeled malate can be reduced to double-labeled αKG, serving as a precursor for m+2 Glu. Meanwhile, citrate synthase combines double-labeled oxaloacetate and single-labeled acetyl-CoA to generate triple-labeled citrate. Subsequent oxidative decarboxylation leads to double-labeled αKG, serving as another source of m+2 Glu. This latter oxidative route can theoretically still explain the observation of m+2 Glu, but not m+3 Glu, in models that do not allow for reductive flux to αKG (Fig. 2 and 6). White circles, ^12^C; black circles, ^13^C; half-filled circles, either ^12^C or ^13^C due to scrambling from the symmetry of fumarate and succinate.

**Fig. S2.**
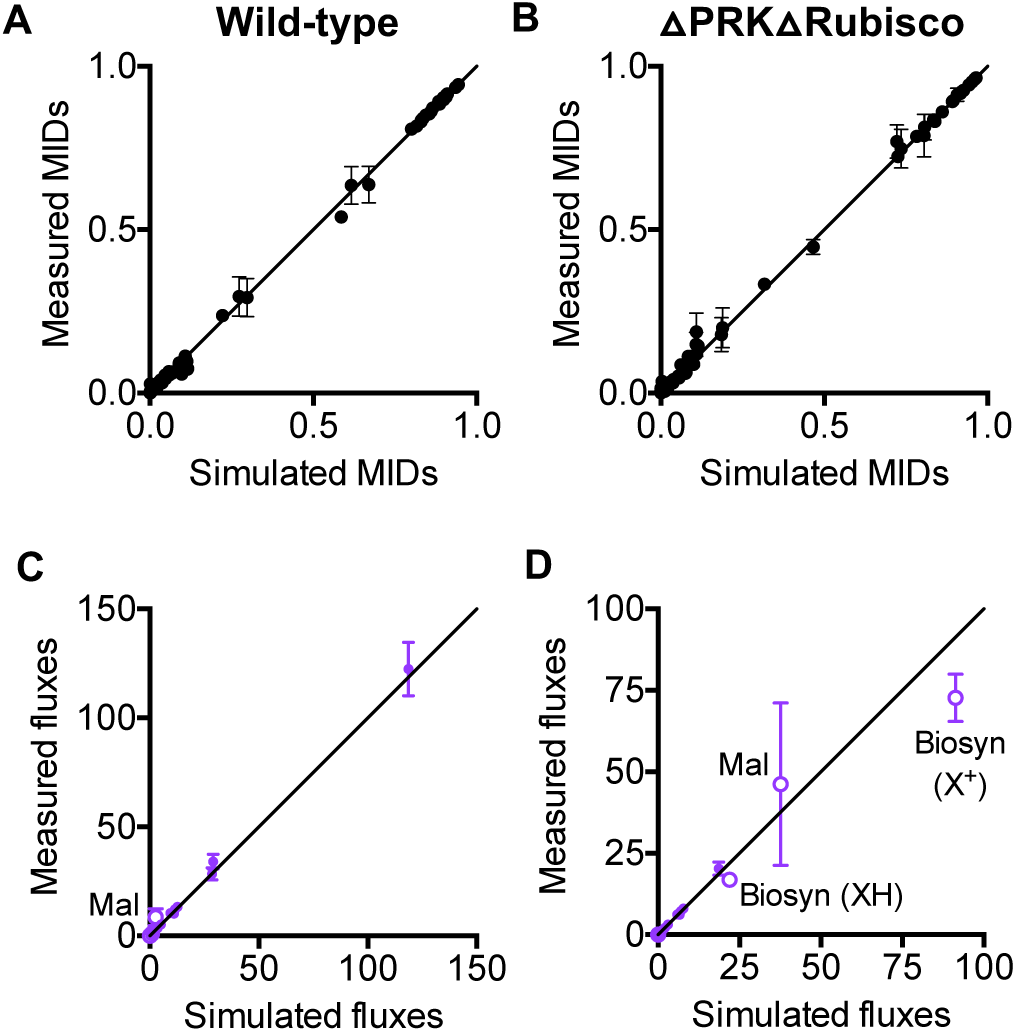
Measured and simulated MIDs and fluxes are similar except for malate excretion and biosynthetic redox reactions for *Rs. rubrum*. Comparison of measured and simulated MIDs (A, B) or flux values (C, D) from optimal flux solutions for WT *Rs. rubrum* (A, C) or the ΔPRKΔRubisco mutant (B, D) grown with [1,4-^13^C]fumarate. (C, D) Values are mole % of the fumarate uptake rate. Error bars, SD; n = 4 for MIDs and malate excretion. Biosynthetic fluxes were assumed to have an SD of 10% of the mean.

**Fig. S3.**
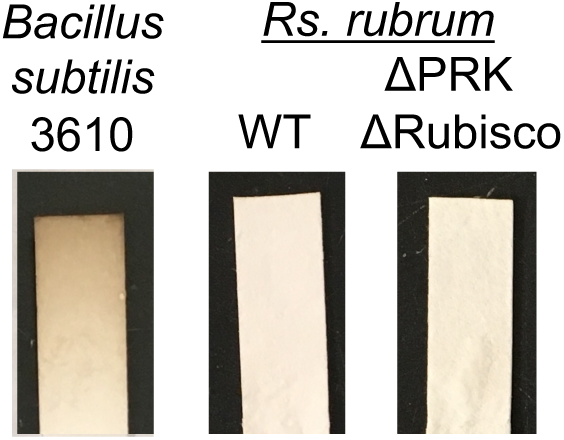
*Rs. rubrum* cultures test negative for sulfide production using lead acetate strips. Strips darken when exposed to sulfide. An aerobic culture of *Bacillus subtilis* 3610 grown in LB medium was used as a positive control for sulfide production (52). Similar trends were observed for two other biological replicates.

**Fig. S4.**
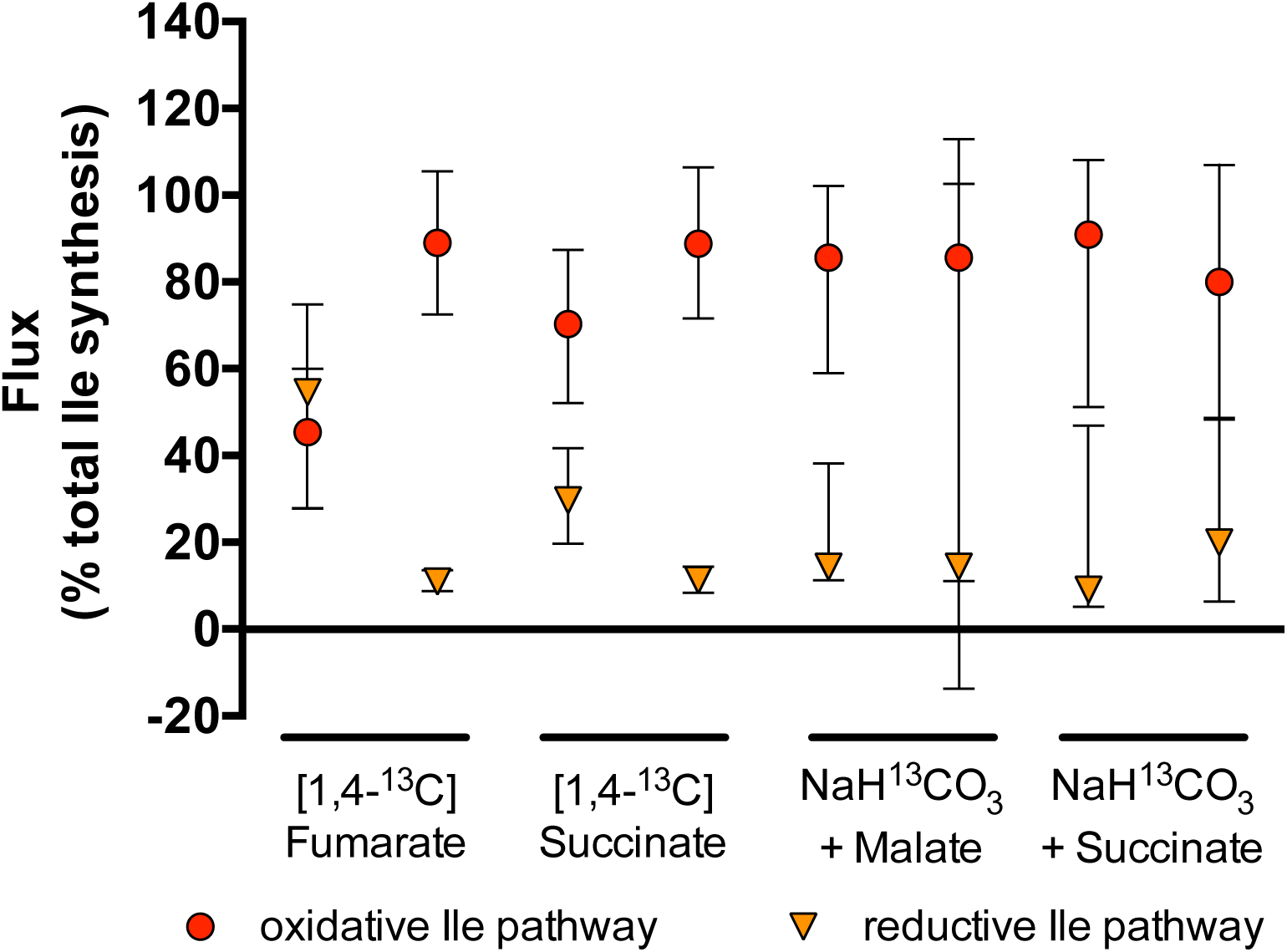
*Rp. palustris* does not adjust fluxes through the reductive and oxidative isoleucine synthesis pathways in response to the absence of the Calvin cycle. Estimated fluxes as a percent of total isoleucine synthesis for the oxidative citramalate-dependent pathway and the reductive threonine-dependent pathway in *Rp. palustris* NifA* (CGA676) and the NifA* ΔPRKΔRubisco strain (CGA4011) during growth on the carbon sources indicated. All flux values are in Table S4.

